## Supplementary information for "Structural basis for membrane attack complex inhibition by CD59"

Figures S1-S9

Table S1

Table S2

Movie S1

**Supplementary Figures**

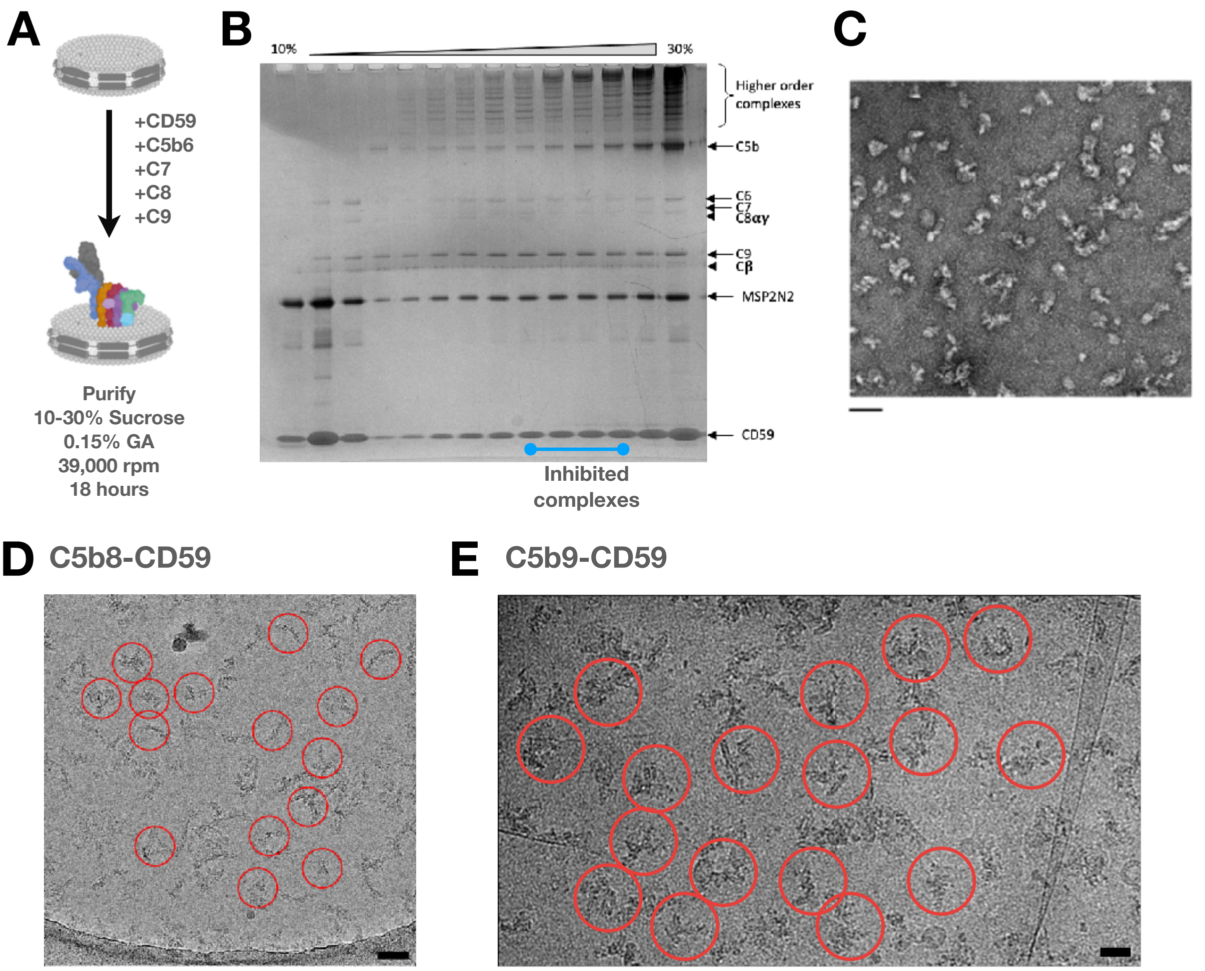

**Figure S1**. Supplementary cryoEM data. (**A**) Schematic showing the assembly and purification of CD59-inhibited complement complexes on lipid nanodiscs. (**B**) Non-reduced SDS-PAGE gel of fractions along the density centrifugation gradient (10-30% sucrose). Sizes corresponding to individual complement proteins, CD59 and MSP2N2 from the nanodisc are indicated. Fractions pooled for further structural analysis are indicated by a blue line. (**C**) Negatively stained C5b9-CD59 complexes corresponding to pooled fractions in (B). Scale bar, 50 nm. (**D**) Representative cryoEM micrograph of C5b8-CD59 sample, individual complexes circled. Scale bar, 40 nm. (**E**) Representative cryoEM micrograph of C5b9-CD59 sample, individual complexes circled. Scale bar, 20 nm.

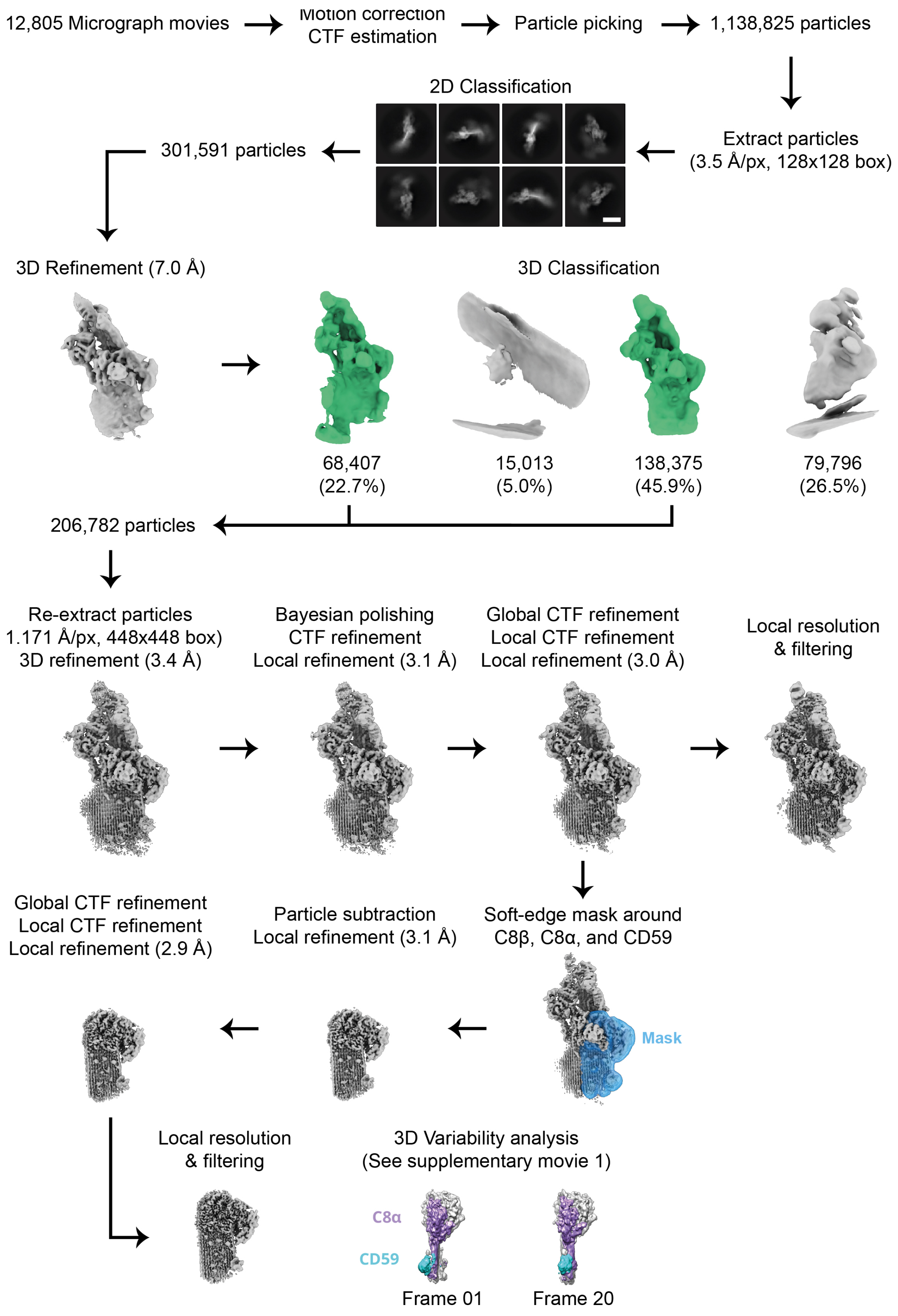

**Figure S2**. CryoEM image processing workflow for C5b8-CD59 map. Schematic outlines steps performed to obtain the structure of the C5b8-CD59 complex. Scale bar referring to the 2D class averages is 13 nm. (see Materials and Methods for details).

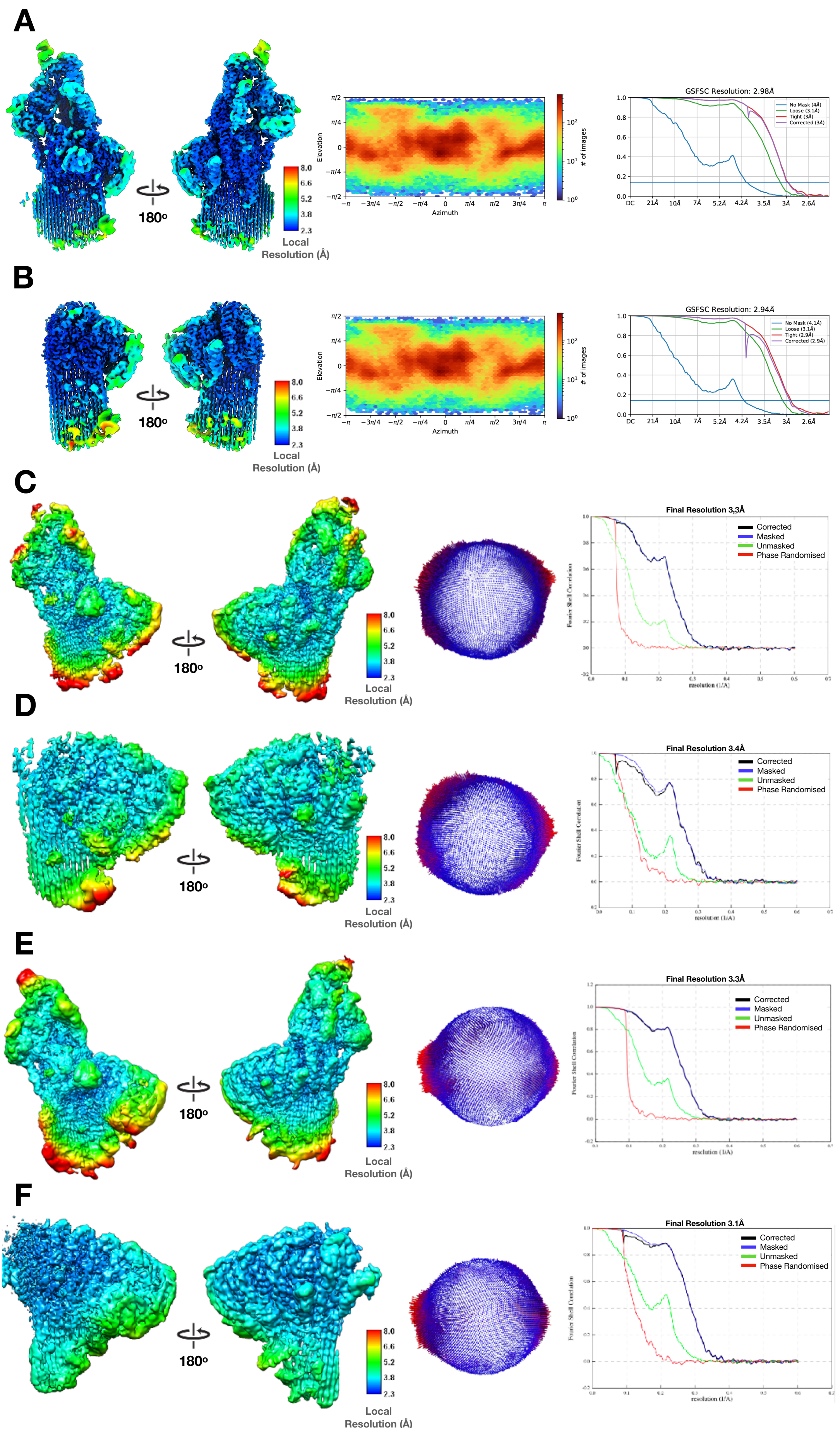
A
180º
Local Resolution (Å)
8.0
6.6
5.2
3.8
2.3
Elevation
Azimuth
### of images
GSFSC Resolution: 2.98Å
No Mask (4Å)
Loose (3.1Å)
Tight (3Å)
Corrected (3Å)
B
180º
Local Resolution (Å)
Elevation
Azimuth
### of images
GSFSC Resolution: 2.94Å
No Mask (4.1Å)
Loose (3.1Å)
Tight (2.9Å)
Corrected (2.9Å)
C
180º
Local Resolution (Å)
Final Resolution 3.3Å
Corrected
Masked
Unmasked
Phase Randomised
D
180º
Local Resolution (Å)
Final Resolution 3.4Å
Corrected
Masked
Unmasked
Phase Randomised
E
180º
Local Resolution (Å)
Final Resolution 3.3Å
Corrected
Masked
Unmasked
Phase Randomised
F
180º
Local Resolution (Å)
Final Resolution 3.1Å
Corrected
Masked
Unmasked
Phase Randomised

**Figure S3**. Map validation information for C5b8-CD59 and C5b9-CD59 complexes. Map validations for C5b8-CD59 (**A**), density subtracted focus refined C5b8-CD59 (**B**), $C5b9_2$-CD59 (**C**), density subtracted focus refined $C55b9_2$-CD59 (**D**), $C5b9_3$-CD59 (**E**), density subtracted focus refined $C5b9_3$-CD59 (**F**). Colored local resolution filtered maps for each reconstruction are shown in the left panels. Angular distribution plots for each reconstruction are shown in the middle panels. FSC curves for each reconstruction are shown in the right panels.

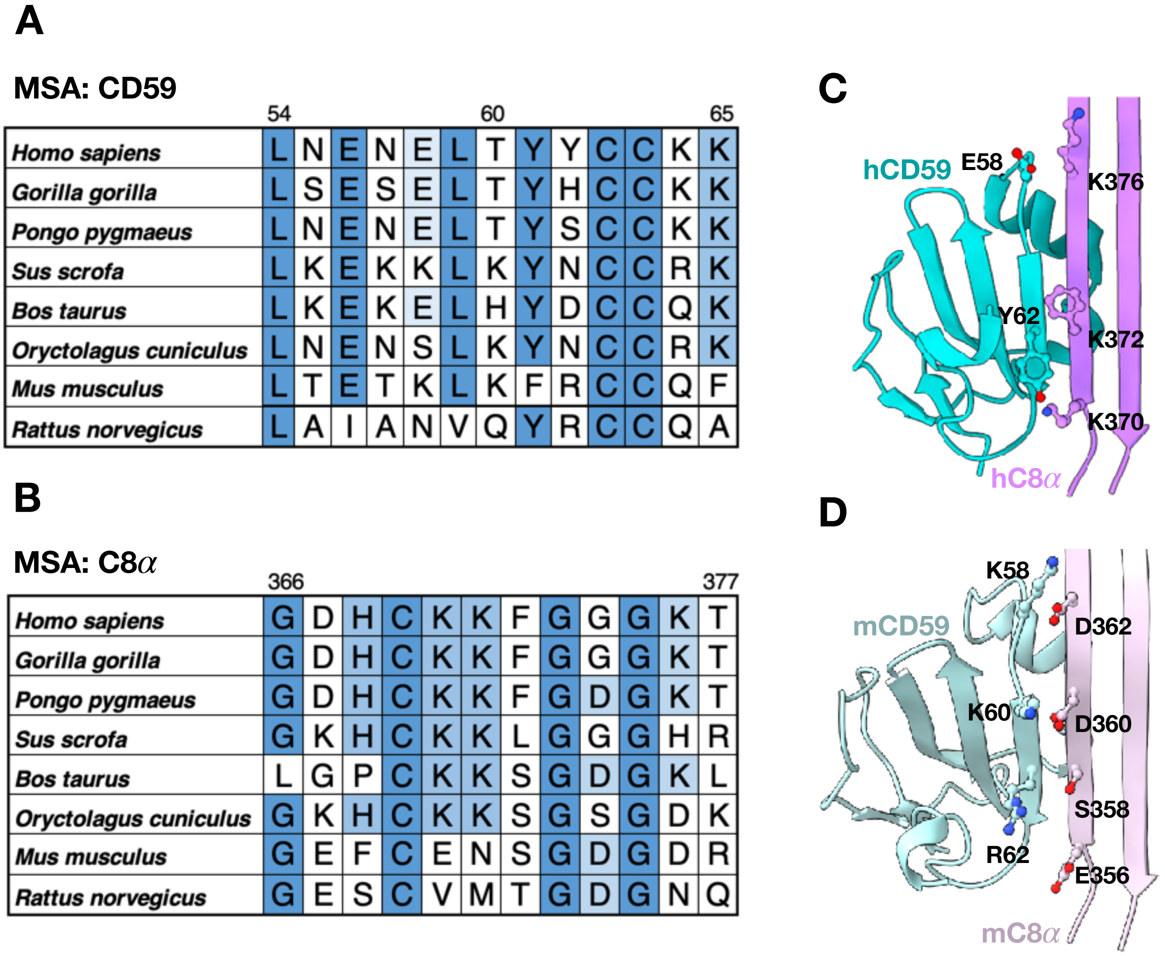

**Figure S4**. Co-evolution analysis of the C8α-CD59 interface. Multisequence alignment (MSA) for CD59 (**A**) and C8α residues (**B**) within the binding interface. Numbering corresponds to the human sequence. (**C**) Ribbon diagram of the human C8α-CD59 interface in our structure. (**D**) Murine model of the interface derived from AlphaFold2 prediction of CD59 (AlphaFold

ID: O55186) and template-based model building of C8α. Key residues that mediate the interactions are shown as sticks.

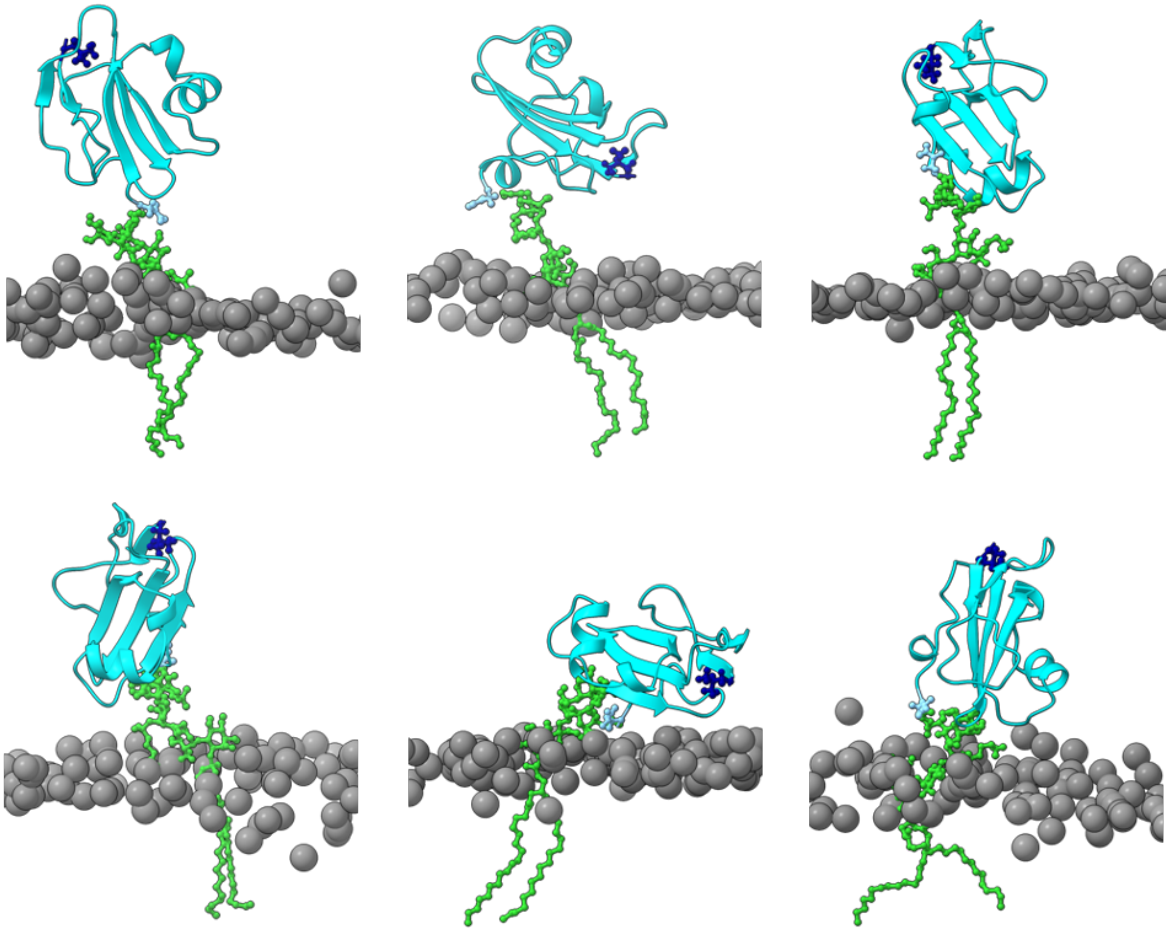

**Figure S5**. MD simulations show flexibility of GPI-anchored CD59. Representative frames sampled from the atomistic MD simulations of GPI-anchored CD59 in a DOPC lipid bilayer. CD59 is shown as cyan ribbons with the GPI anchor in green sticks and the N- and C- terminal residues in dark blue and light blue sticks, respectively. Phosphorous atoms from lipid headgroups are grey spheres.

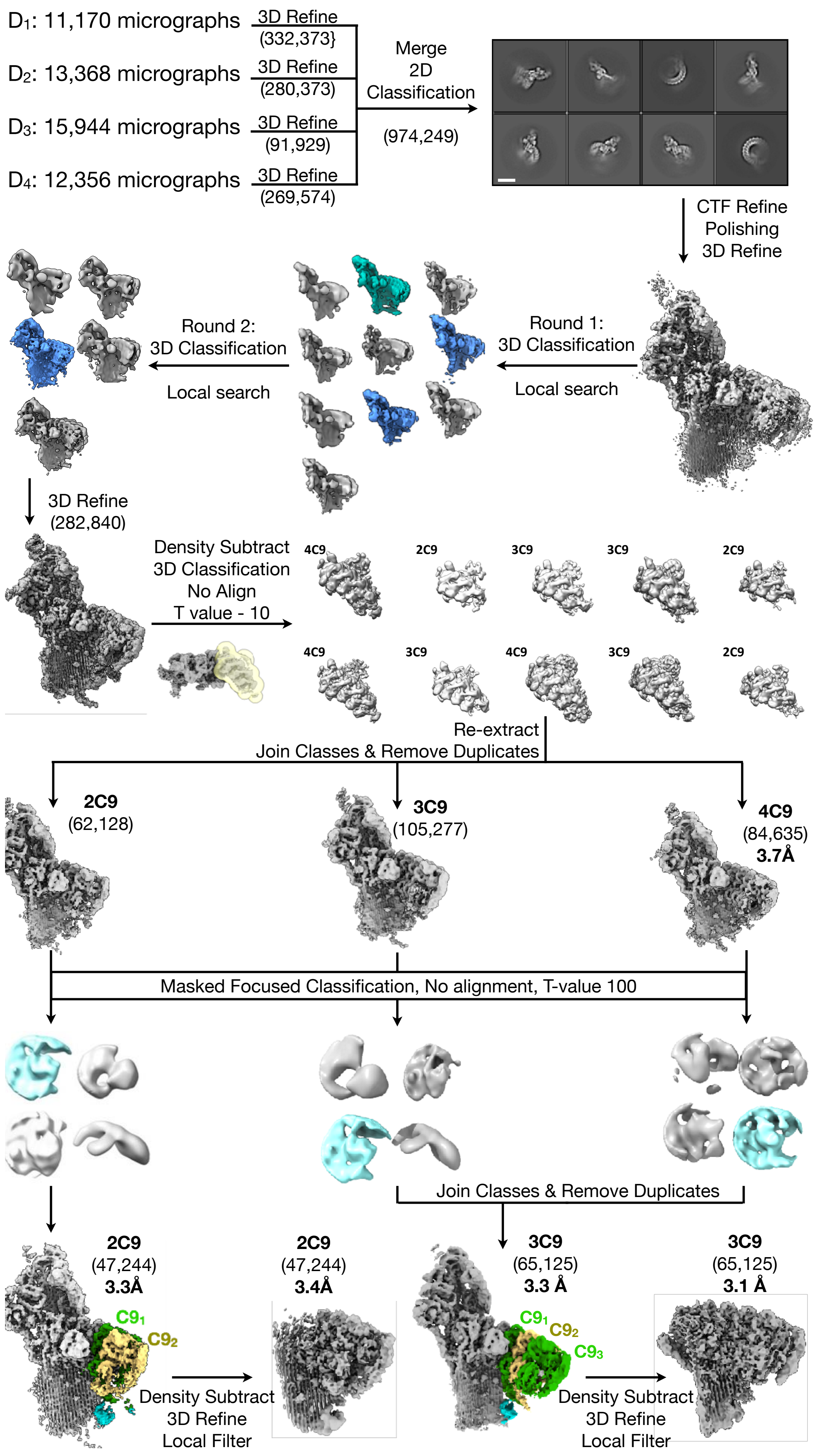

D1: 11,170 micrographs
D2: 13,368 micrographs
D3: 15,944 micrographs
D4: 12,356 micrographs
3D Refine (332,373}
3D Refine (280,373)
3D Refine (91,929)
3D Refine (269,574)
Merge 2D Classification
(974,249)
CTF Refine Polishing 3D Refine
Round 1: 3D Classification
Local search
Round 2: 3D Classification
Local search
3D Refine (282,840)
Density Subtract 3D Classification No Align T value - 10
4C9 2C9 3C9 3C9 2C9
4C9 3C9 4C9 3C9 2C9
Re-extract
Join Classes & Remove Duplicates
2C9 (62,128)
3C9 (105,277)
4C9 (84,635) 3.7Å
Masked Focused Classification, No alignment, T-value 100
Join Classes & Remove Duplicates
2C9 (47,244) 3.3Å
C9₁ C9₂
Density Subtract 3D Refine Local Filter
2C9 (47,244) 3.4Å
3C9 (65,125) 3.3 Å
C9₁ C9₂ C9₃
Density Subtract 3D Refine Local Filter
3C9 (65,125) 3.1 Å

**Figure S6**. CryoEM image processing workflow for C5b9-CD59. Schematic outlines steps performed to obtain the $C5b9_2$-CD59 and $C5b9_3$-CD59 structures. Scale bar referring to the 2D class averages is 20 nm. Particle numbers are in brackets. (see Materials and Methods for details).

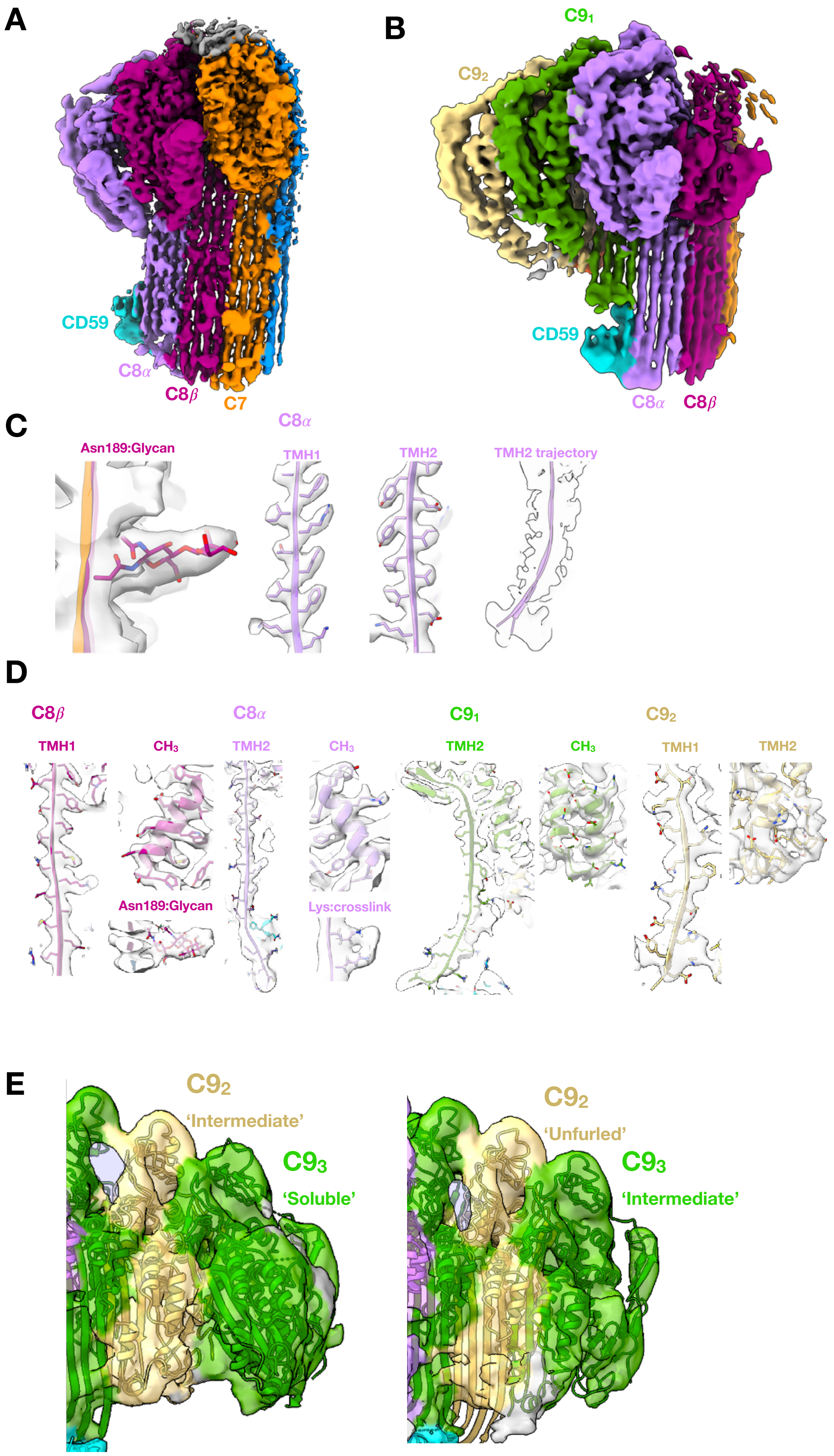

A
CD59
C8α
C8β
C7
B
C9₁
C9₂
CD59
C8α
C8β
C
C8α
Asn189:Glycan
TMH1
TMH2
TMH2 trajectory
D
C8β
TMH1
CH₃
Asn189:Glycan
C8α
TMH2
CH₃
Lys:crosslink
C9₁
TMH2
CH₃
C9₂
TMH1
TMH2
E
C9₂
'Intermediate'
C9₃
'Soluble'
C9₂
'Unfurled'
C9₃
'Intermediate'

**Figure S7**. Map quality of the C5b8-CD59 and C5b9-Cd59 structures. (**A-B**) Density subtracted focus refined maps for the C5b8-CD59 (**A**) and C5b9$_2$-CD59 (**B**) complexes. (**C-D**) Fits of the C5b8-CD59 (**C**) and C5b9$_2$-CD59 (**D**) models into their respective density subtracted maps. (**E**) Models for two conformations of the terminal C9 (C9$_3$) derived from the cyroDRGN analysis of the C5b9-CD59 complex.

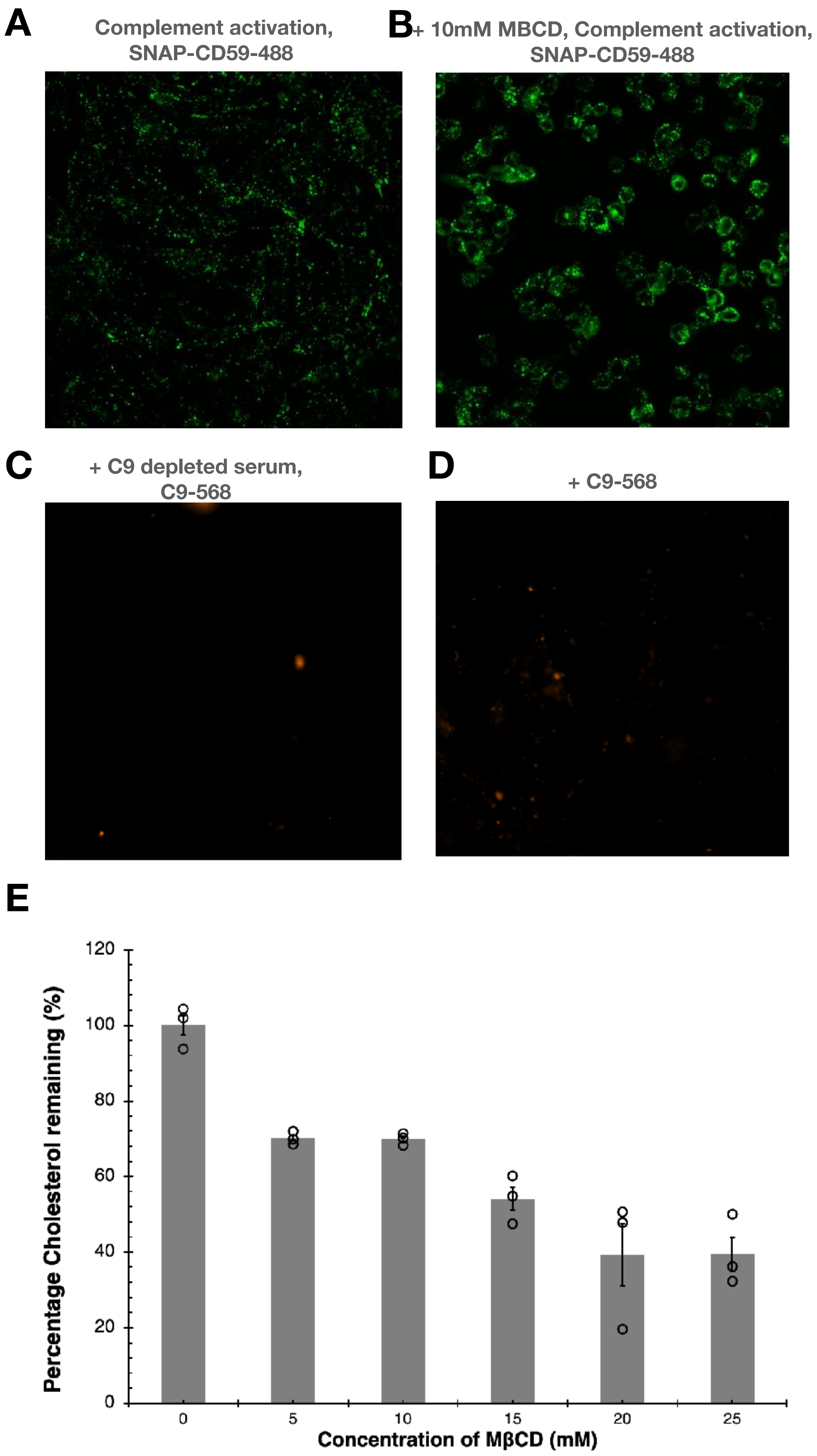

A
Complement activation,
SNAP-CD59-488
B
+ 10mM MBCD, Complement activation,
SNAP-CD59-488
C
+ C9 depleted serum,
C9-568
D
+ C9-568
E
Percentage Cholesterol remaining (%)
120
100
80
60
40
20
0
0
5
10
15
20
25
Concentration of MβCD (mM)

**Figure S8**. Supporting cellular assay controls. (**A-B**) CHO cells expressing SNAP-CD59 were treated with a polyclonal anti-CHO IgG antibody to activate complement. Cells were incubated with C9-delpeted human serum supplemented with a chemically-labeled fluorescent C9 (C9-Alexafluor 568) capable of forming MAC. CD59 was visualized with SNAP-Oregon (488 nm) in control cells (**A**) and in cells treated with MβCD to deplete cholesterol (**B**). Wide-field fluorescence microscopy was used to visualize C9 on the cell surface in the absence of complement activator (**C**) and when incubated with C9-Alexafluor 568 alone (**D**). No nonspecific binding is observed. (**E**) Amplex-Red cholesterol depletion assay detecting the extent of cholesterol present in the cells normalized to untreated cells. Individual measurements are given as points, the average and standard deviation across three technical replicates are shown.

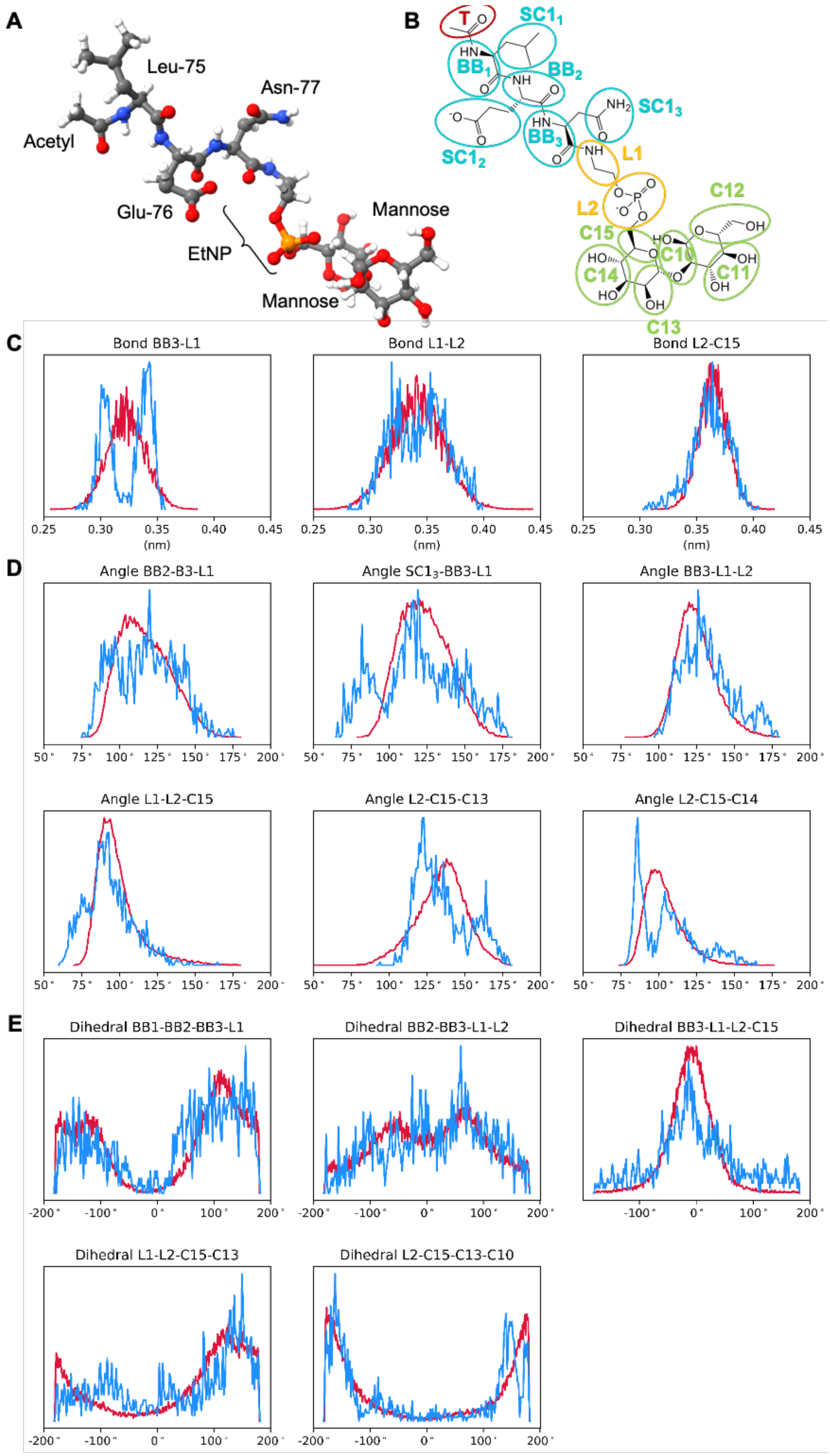
A
Leu-75
Asn-77
Acetyl
Glu-76
EtNP
Mannose
Mannose
B
T
SC1₁
BB₁
BB₂
SC1₃
SC1₂
BB₃
L1
L2
C12
C15
C10
C14
C11
C13
C
Bond BB3-L1
Bond L1-L2
Bond L2-C15
(nm)
D
Angle BB2-B3-L1
Angle SC1₃-BB3-L1
Angle BB3-L1-L2
Angle L1-L2-C15
Angle L2-C15-C13
Angle L2-C15-C14
E
Dihedral BB1-BB2-BB3-L1
Dihedral BB2-BB3-L1-L2
Dihedral BB3-L1-L2-C15
Dihedral L1-L2-C15-C13
Dihedral L2-C15-C13-C10

**Figure S9**. Parameterization of the phosphoethanolamine linker (EtNP). (**A**) Stick representation of the molecule used for parameterization. (**B**) Coarse-grained mapping of the molecule in (A). Beads corresponding to the CD59 residues are in cyan, the terminal acetyl cap is red, beads for the EtNP linker (L1 and L2) are in yellow, and beads corresponding to the mannose residues are in green. Comparing bond lengths (**C**) angles (**D**) and dihedral angles (**E**) involving beads L1 and L2 of the EtNP linker between atomistic (blue) and coarse-grained (red) simulations.

**Table S1**. Validation statistics for EM maps and models.

| | **#1 C5b8-CD59**<br>EMD: | **#2 C5b8-CD59$^{arc}$**<br>EMD: | **#3 C5b9$_2$-CD59**<br>EMD: | **#4 C5b9$_3$-CD59**<br>EMD: | **#5 C5b9$_2$-CD59$^{arc}$**<br>EMD: | **#6 C5b9$_3$-CD59$^{arc}$**<br>EMD: |
|---|---|---|---|---|---|---|
| **Data collection and processing** | | | | | | |
| **Magnification** | 105k | 105k | 105k | 105k | 105k | 105k |
| **Voltage (kV)** | 300 | 300 | 300 | 300 | 300 | 300 |
| **Electron exposure (e–/Å$^2$)** | 50 | 50 | $D_1$ 43<br>$D_2$ 56<br>$D_3$ 40<br>$D_4$ 50 | $D_1$ 43<br>$D_2$ 56<br>$D_3$ 40<br>$D_4$ 50 | $D_1$ 43<br>$D_2$ 56<br>$D_3$ 40<br>$D_4$ 50 | $D_1$ 43<br>$D_2$ 56<br>$D_3$ 40<br>$D_4$ 50 |
| **Defocus range (μm)** | -1.0 to –2.25 | -1.0 to –2.25 | -1.0 to -2.25 | -1.0 to -2.25 | -1.0 to -2.25 | -1.0 to -2.25 |
| **Detector Type** | Falcon IV | | K3 | K3 | K3 | K3 |
| **Pixel size (Å)** | 1.171 | 1.171 | $D_1$ 0.829<br>$D_2$ 0.831<br>$D_3$ 0.831<br>$D_4$ 0.85 | $D_1$ 0.829<br>$D_2$ 0.831<br>$D_3$ 0.831<br>$D_4$ 0.85 | $D_1$ 0.829<br>$D_2$ 0.831<br>$D_3$ 0.831<br>$D_4$ 0.85 | $D_1$ 0.829<br>$D_2$ 0.831<br>$D_3$ 0.831<br>$D_4$ 0.85 |
| **Symmetry imposed** | C1 | C1 | C1 | C1 | C1 | C1 |
| **Initial particle images (no.)** | 1,138,825 | 1,138,825 | $D_1$ 737,138<br>$D_2$ 1,058,026<br>$D_3$ 1,330,232<br>$D_4$ 722,870 | $D_1$ 737,138<br>$D_2$ 1,058,026<br>$D_3$ 1,330,232<br>$D_4$ 722,870 | $D_1$ 737,138<br>$D_2$ 1,058,026<br>$D_3$ 1,330,232<br>$D_4$ 722,870 | $D_1$ 737,138<br>$D_2$ 1,058,026<br>$D_3$ 1,330,232<br>$D_4$ 722,870 |
| **Particle images before merging (no.)** | N/A | N/A | $D_1$ 332,373<br>$D_2$ 280,769<br>$D_3$ 91,929<br>$D_4$ 269,574 | $D_1$ 332,373<br>$D_2$ 280,769<br>$D_3$ 91,929<br>$D_4$ 269,574 | $D_1$ 332,373<br>$D_2$ 280,769<br>$D_3$ 91,929<br>$D_4$ 269,574 | $D_1$ 332,373<br>$D_2$ 280,769<br>$D_3$ 91,929<br>$D_4$ 269,574 |
| **Final particle images (no.)** | 206,782 | 206,782 | 47,244 | 33,138 | 47,244 | 33,138 |
| **Map resolution (Å)**<br>FSC threshold 0.143 | 3.0 | 2.9 | 3.3 | 3.3 | 3.5 | 3.2 |
| **Map resolution range (Å)** | 2.3-8.0 | 2.3-8.0 | 3.0-12.4 | 3.0-10.2 | 3.1-11.0 | 3.0-8.0 |
| **Refinement** | | | | | | |
| **Initial model used (PDB code)** | 7NYD, 2J8B | 7NYD,2J8B | 7NYD, 2J8B | 7NYD, 2J8B | 7NYD, 2J8B | 7NYD, 2J8B |
| **Model resolution (Å)** | | | | | | |
| FSC threshold 0.5 | 3.0 | | 3.3 | 3.2 | | |
| **Map sharpening**<br>*B* factor (Å$^2$) | -60 | -60 | -50 | -50 | -50 | -50 |
| **Model composition** | | | | | | |
| Non-hydrogen atoms | 30,943 | | 37,288 | 40,505 | | |
| Protein residues | 3910 | | 4704 | 5113 | | |
| Ligands | BMA: 1 | | BMA: 4 | BMA: 0 | | |
| Ions | NAG: 6 | | NAG: 8 | NAG: 10 | | |
| | Ca: 2 | | 0 | 0 | | |
| ***B* factors (Å$^2$)** | | | | | | |
| **Protein** | 86.80 | | 117.93 | 118.03 | | |
| **Ligand** | 136.15 | | 187.03 | 138.05 | | |
| **R.m.s. deviations** | | | | | | |
| Bond lengths (Å) | 0.003 | | 0.001 | 0.009 | | |
| Bond angles (°) | 0.739 | | 0.867 | 0.791 | | |
| **Validation** | | | | | | |
| MolProbity score | 1.81 | | 1.87 | 1.66 | | |
| Clashscore | 4.43 | | 8.53 | 5.49 | | |
| Poor rotamers (%) | 1.4 | | 0.56 | 0.31 | | |
| **Ramachandran plot** | | | | | | |
| Favored (%) | 92.24 | | 95.87 | 94.80 | | |
| Allowed (%) | 7.76 | | 5.13 | 5.20 | | |
| Disallowed (%) | 0.0 | | 0.0 | 0.0 | | |

**Table S2**. CG parameters of the molecules used for parametrization of the EtNP linker.

| Bead name | Bead type | Bond | $r_0$ (nm) | $K_b$ (kJ/mol) | Angle | $\theta_0$ (°) | $K_a$ (kJ/mol) |
|---|---|---|---|---|---|---|---|
| T | SN0 | T – $BB_1$ | 0.3 | 20000 | T – $BB_1$ – $BB_2$ | 128 | 20.0 |
| $BB_1$ | P5 | $BB_1$ – $SC1_1$ | 0.33 | 7500 | T – $BB_1$ – $SC1_1$ | 108 | 30.0 |
| $SC1_1$ | C1 | $BB_1$ – $BB_2$ | 0.39 | 20000 | $BB_1$ – $BB_2$ – $BB_3$ | 132 | 30.0 |
| $BB_2$ | P5 | $BB_2$ – $SC1_2$ | 0.4 | 5000 | $SC1_2$ – $BB_2$ – $BB_3$ | 108 | 30.0 |
| $SC1_2$ | Qa | $BB_2$ – $BB_3$ | 0.35 | 20000 | $BB_2$ – $BB_3$ – $SC1_3$ | 96 | 80.0 |
| $BB_3$ | P5 | $BB_3$ – $SC1_3$ | 0.32 | 5000 | $BB_2$ – $BB_3$ – L1 | 105 | 2.5 |
| $SC1_3$ | P5 | $BB_3$ – L1 | 0.33 | 10000 | $SC1_3$ – $BB_3$ – L1 | 90 | 1.0 |
| L1 | GSNda | L1 – L2 | 0.35 | 5000 | $BB_3$ – L1 – L2 | 130 | 60.0 |
| L2 | GQa | L2 – C15 | 0.365 | 18000 | L1 – L2 – C15 | 45 | 5.0 |
| C15 | GNa | C15 – C14 | 0.33 | 20000 | L2 – C15 – C13 | 180 | 15.0 |
| C14 | GP3 | C15 – C13 | 0.35 | 20000 | L2 – C15 – C14 | 70 | 20.0 |
| C13 | GSP1 | C14 – C13 | 0.28 | 40000 | C15 – C13 – C10 | 65 | 45.0 |
| C12 | GP2 | C13 – C10 | 0.36 | 20000 | C14 – C13 – C10 | 95 | 100.0 |
| C11 | GP3 | C12 – C11 | 0.33 | 30000 | | | |
| C10 | GSN0 | C12 – C10 | 0.35 | 30000 | | | |
| | | C11 – C10 | 0.28 | 40000 | | | |

| Dihedral | Type | $\phi_0$ (°) | $K_d$ (kJ/mol) |
|---|---|---|---|
| $BB_2$ – $BB_3$ – L1 – L2 | Proper | 220 | 1.5 |
| $BB_3$ – L1 – L2 – C15 | Proper | –195 | 4.0 |
| L1 – L2 – C15 – C13 | Proper | 320 | 3.5 |
| L2 – C15 – C13 – C10 | Proper | 30 | 3.0 |
| $BB_1$ – $BB_2$ – $BB_3$ – L1 | Improper | –200 | 3.0 |

**Movie S1**. 3D variability analysis of the focus refined C5b8-CD59 map. A linear movie of volumes visualizing variability across the first principal component is shown. Volumes are colored according to protein components: CD59 is cyan, C8α is pink, all other proteins in the map are grey.
